# *SLTAB2* is the paramutated *SULFUREA* locus in tomato

**DOI:** 10.1101/033605

**Authors:** Quentin Gouil, Ondřej Novák, David Baulcombe

## Abstract

The *sulfurea (sulf)* allele is a silent epigenetic variant of a tomato gene (*Solanum lycopersicum)* affecting pigment production. It is homozygous lethal but, in a heterozygote *sulf*/+, the wild type allele undergoes silencing so that the plants exhibit chlorotic sectors. This transfer of the silenced state between alleles resembles the process of paramutation that is best characterised in maize. To understand the mechanism of paramutation we mapped *SULF* to the ortholog *SLTAB2* of an *Arabidopsis* gene that, consistent with the pigment deficiency, is involved in the translation of photosystem I. Paramutation of *SLTAB2* is linked to an increase in DNA methylation and production of small interfering RNAs at its promoter. Virus-induced gene silencing of *SLTAB2* phenocopies *sulf* consistent with the possibility that siRNAs mediate the paramutation of *SULFUREA*. Unlike the maize systems the paramutagenicity of *sulf* is not, however, associated with repeated sequences at the region of siRNA production or DNA methylation.

## Introduction

Paramutation involves transfer of an epigenetic mark from a (paramutagenic) silent gene to the active (paramutable) allele so that it becomes heritably paramutagenic and silent. Several plant species exhibit paramutation and the best characterised examples, the *b1* and *pl1* loci in maize, have been linked to the process of RNA-directed DNA methylation (RdDM)(Chandler and Stam 2004; Hollick 2012) in which paramutagenic small interfering (si)RNAs mediate silencing of the paramutable allele. This simple model does not, however, explain why most siRNA loci are not paramutagenic: there must be other factors.

To shed light on the mechanism of paramutation we are analysing the tomato *SULFUREA (SULF)* locus. The silent *sulf* allele has a chlorotic phenotype (Hagemann 1958), reduced auxin (Ehlert, Schottler, Tischendorf, Ludwig-Muller, and Bock 2008) and it is homozygous lethal. A heterozygous *sulf*/+ plant is viable but it has large chlorotic sectors that are due to paramutation of the active allele to a silenced state in early development. This paramutated state is heritable and paramutagenic (Hagemann 1969). *SULF* maps to the pericentromeric heterochromatin of chromosome 2, at approximately 29 cM from the *S* locus (Solyc02g077390, compound inflorescence)(Hagemann and Snoad 1971) but the affected gene could not be mapped precisely due to low recombination frequency (The Tomato Genome Consortium 2012).

A gene ortholog of the *Arabidopsis ATAB2* is strongly down-regulated in chlorotic sectors of *sulf*/+ tomato (Ehlert, Schottler, Tischendorf, Ludwig-Muller, and Bock 2008) but it was previously excluded as the *SULF* gene because it is still expressed at detectable levels. However, from analysis of transcriptome, methylome and small RNA populations of wild-type and paramutated tomato leaves we show here that paramutation of *SLTAB2* is responsible for the *sulf* chlorosis and decrease in auxin levels. *SLTAB2* silencing was associated with changes in DNA methylation and siRNA levels at its promoter, a signature of RdDM.

Additional evidence supporting the identification of *SLTAB2* as *SULF* is from virus-induced gene silencing (VIGS) of the *SLTAB2* promoter resulting in the methylation of the target DNA sequence, silencing of its expression and a phenocopy of the *sulf* chlorosis. Together these results support a causal role of siRNAs and RdDM in paramutation but, unlike the maize examples, the *SLTAB2/SULF* locus lacked repeated sequences. Mapping of *SULF* to *SLTAB2* and further comparison with maize will help build a general model of paramutation in plants.

## Materials and Methods

### Plant material and growth conditions

*Atab2* T-DNA knockouts (GABI-KAT line 354B01) and wild-type Col-0 seedlings were sown on 1/2 MS, 1x Nitsch & Nitsch vitamins, 0.8% agar, 1.5% sucrose, pH 6; stratified for 72 h at 4°C in the dark and transferred to short-day conditions (8 h light at 23°C and 50 *μ*mol photons m^2^.s^1^, 16 h dark at 21°C). Whole seedlings were collected after seven days of growth. Tomato plants were raised from seeds in compost (Levington M3) and maintained in a growth room at 23°C with 16-h light and 8-h dark periods with 60% relative humidity, at a light intensity of 150 *μ*mol photons m^−2^.s^−1^. Young leaves were collected from one-month-old plants. *Sulf* and *sulf* /+ tissue was collected from *sulf* /+ plants that had both fully yellow (*sulf*) and fully green (*sulf*/+) sectors.

### Transcriptome analysis

Total RNA samples were prepared from 100 mg of leaf tissue using TRIzol (Life Technologies). For qPCR, 5 *μ*g of total RNA was first DNase treated using Turbo DNase (Ambion), following the manufacturers guidelines. cDNA was then synthesised using random hexamers and oligodT and SuperScript III (Life Technologies), according to the protocol. qPCR was performed on a Roche LC480 with SYBR in technical triplicates. mRNA abundance was normalised by the geometric mean of two housekeeping genes *TIP41* and *EXPRESSED* (Coker and Davies 2003). Genotyping of amplified cDNA was performed by digesting 100 ng purified *SLTAB2* amplicon with BaeI (NEB) for 12 hours at 25°C in 1x NEB2.1, 100 *μ*g.ml^−1^ BSA, 20 *μ*M SAM as per manufacturer’s instructions, and electrophoresis on 1.5% agarose gel. Strand-specific RNA-Seq libraries for two wild-types and three pairs of *sulf* and *sulf* /+ were made and indexed with the ScriptSeq v2 kit (Epicentre) according to the protocol after RiboZero treatment (Plant leaf, Epicentre), and sequenced as a pool on one lane of HiSeq2000 100PE. Sequences were trimmed and filtered with Trim Galore! with default parameters and 11–29 millions reads per library were concordantly aligned on Heinz genome SL2.50 and ITAG2.4 gene models using TopHat2 v2.0.13 (with parameters -r 200 --mate-std-dev 100 -N 3 --read-edit-dist 3 --library-type fr-firststrand --solexa1.3-quals, and version 2.2.4.0 of Bowtie). Differential expression analysis was performed on raw counts on annotated mRNAs (ITAG2.4) with DESeq2 v1.8.1. Genes were considered differentially expressed when the adjusted p-value was < 0.05. Hierarchical correlation clustering of the genes differentially expressed between wt and *sulf* was performed in SeqMonk (v0.32.0). Gene Ontology analysis was performed with the goseq package (v1.20.0) using previously published gene ontology annotation (Chitwood, Maloof, and Sinha 2013), normalising with mRNA length and running with the following parameters: method = Wallenius, repcnt = 2000, use_genes_without_cat = F. Categories were considered to be over-represented if the associated p-value was < 0.05 after Benjamini-Hochberg correction.

### sRNA-Seq

sRNAs were cloned from 10 *μ*g total RNA (from the same tissue as used for RNA-Seq, for two wild-types and two pairs of *sulf* and *sulf* /+) using the Illumina TruSeq Small RNA cloning kit and libraries were indexed during the PCR step (12 cycles) according to the manufacturers protocol. Gel size-selected, pooled libraries were sequenced on a HiSeq2000 50SE. Sequences were trimmed and filtered with Trim Galore! (with the adapter parameter -a TGGAATTCTCGGGT-GCCAAGG) and 14–20 million reads per library were mapped without mismatches and clustered on Heinz genome SL2.50 using the ShortStack software v2.1.0 (Axtell 2013). sRNA counts on the defined loci were analysed with DESeq2 v1.8.1. Uniquely mapping reads on DMR1 and DMR2 were normalised with edgeR’s implementation of TMM size factors, on all sRNAs present in all libraries and with at least 10 total counts (Robinson, McCarthy, and Smyth 2010), and a Poisson regression was applied to the normalised counts (generalised linear model in R, with the variable ‘genotype’ taking values wt, *sulf* /+ and *sulf*).

### Methylome analysis

DNA was extracted from 100 mg of leaf tissue (from the same sampling as for RNA-Seq and sRNA-Seq, for two wild-types and two sulf) using the Puregene kit (QIAGEN). Bisulfite library preparation was performed with a custom protocol similar to Urich *et al*., 2015 (Urich, Nery, Lister, Schmitz, and Ecker 2015). 1.2 *μ*g DNA was sonicated on a Covaris E220 to a target size of 400 bp and purified on XP beads (Ampure, ratio 1.8). DNA was end-repaired and A-tailed using T4 DNA polymerase and Klenow Fragment (NEB) and purified again using XP beads (ratio 1.8x). Methylated Illumina Y-shaped adapters for paired-end sequencing were ligated using Quick-Stick Ligase (Bioline). 450 ng of purified (ratio 1.8x), adapter-ligated DNA was bisulfite-converted using EZ DNA Methylation-Gold Kit (Zymo Research) according to manufacturer’s instructions. DNA was barcoded using 12 cycles of PCR amplification with KAPA HiFi HotStart Uracil+ Ready Mix (Kapabiosystems) with PE1.0 and custom index primers (courtesy of the Sanger Institute). Pooled libraries were sequenced to a depth of about 5x per strand on a HiSeq2500 125PE. Sequences were trimmed and filtered with Trim Galore!, then mapped on Heinz genome SL2.50 using Bismark v0.14.3 (Krueger and Andrews 2011) (first in paired-end mode with options --score-min L,0,-0.2 -p 4 --reorder --ignore-quals --no-mixed --no-discordant --maxins 1500-X 1500 --unmapped --ambiguous, then unmapped read1 was mapped in single-end mode with the same quality parameter -N 1). Reads were deduplicated with bismark_deduplicate and methylation calls were extracted using Bismark methylation_extractor (with options -r2 2 for paired end reads). Methylated and unmethylated counts for cytosines of both strands were pooled into contiguous 200 bp bins according to context (CG, CHG and CHH) with a custom python script. Bins with less than 10 counts were excluded from analysis. Bins are considered differentially methylated if the maximum p-value of the two chi-square tests (wt1 vs *sulf* 1, wt2 vs *sulf*2) is < 0.01. Analysis of methylation by McrBC was performed as previously described (Bond and Baulcombe 2015).

### VIGS

DMR1a (606 bp) and DMR1b (562 bp) genomic inserts were cloned into the binary TRV RNA2 vector using the KpnI and XhoI restriction sites of the multiple cloning site as described previously (Liu, Schiff, and Dinesh-Kumar 2002; Bond and Baulcombe 2015). Cotyledons of tomato seedlings were agro-infiltrated 10 days after sowing with a 1:1 mixture of *Agrobacterium tumefaciens* (strain GV3101:pMP90 + pSOUP) carrying TRV RNA1 andRNA2at *OD*_600_ = 1.5. Symptoms of *SLTAB*2 silencing were visible from 2 weeks post-infection.

### Auxin quantification

Endogenous levels of free IAA were detected by LC-MS/MS method as described in (Novák, Hényková, Sairanen, Kowalczyk, Pospíšil, and Ljung 2012). Briefly, 10–20 mg fresh tissue of the control and mutant lines were collected, extracted in ice-cold 50 mM sodium phosphate buffer (pH 7) and purified by SPE on hydrophilic-lipophilic balance reversed-phase sorbent columns (Oasis HLB, 1 cc/30 mg, Waters). To each extract, 5 pmol of ^13^C_6_-IAA were added as internal standards to validate the quantification. Purified samples were analysed by the LC-MS/MS system consisting of an ACQUITY UPLC System (Waters, Milford, MA, USA) and Xevo TQ-S (Waters) triple quadrupole mass spectrometer. Quantification was obtained using a multiple reaction monitoring (MRM) mode of selected precursor ions and the appropriate product ion.

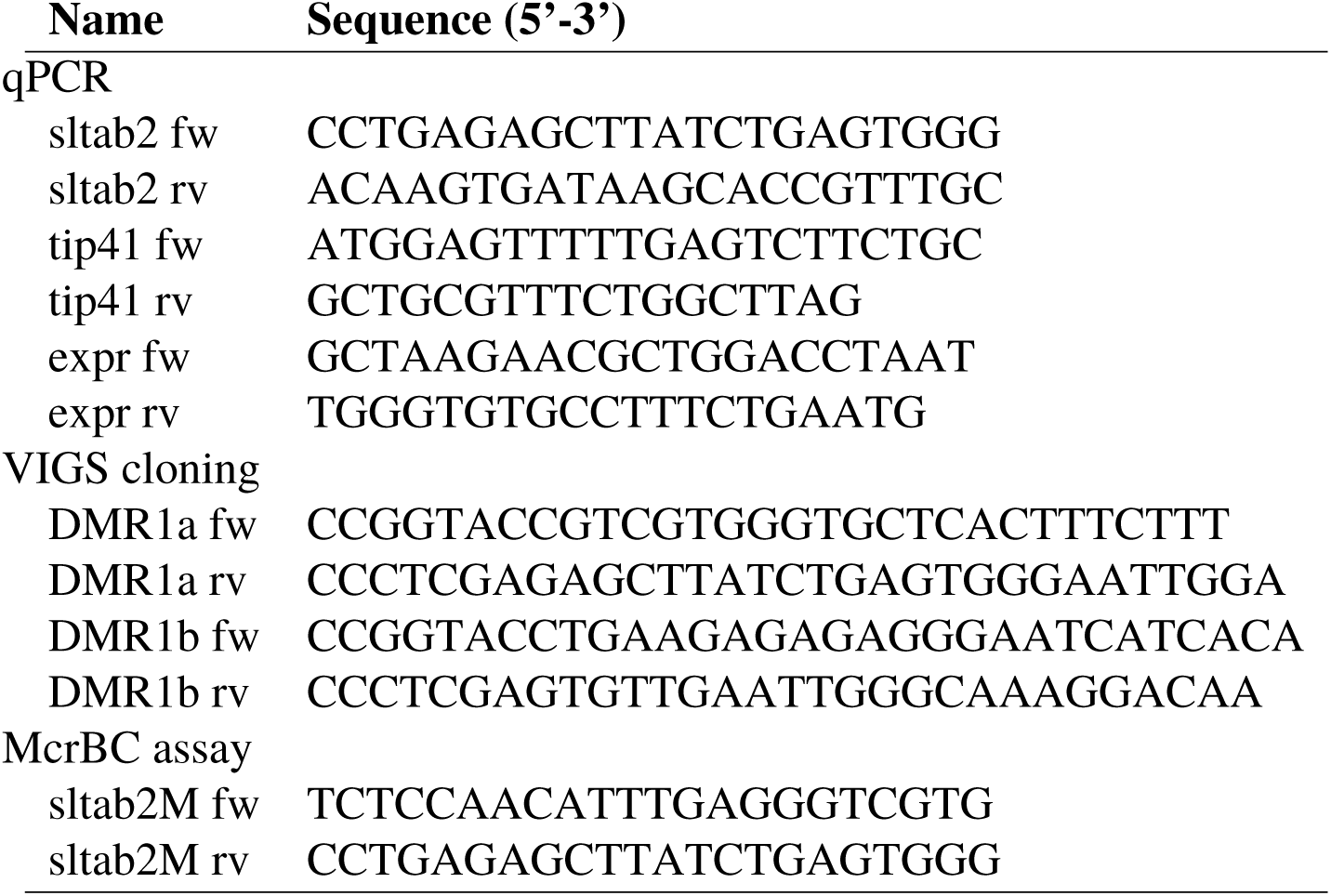
Oligonucleotides used in the present study.

## Oligonucleotides

### Accession codes

All sequencing data have been deposited in the Sequence Read Archive under the BioProject SRP066362.

## Results

### Pericentromeric *SLTAB2* is strongly down-regulated in *sulfurea*

The tomato lines in this study had either unsilenced *SULF* loci (wild-type) or they were the progeny of a cross between wild-type and a plant with sectors with silent *sulf*. Some of the F1 plants were wild-type and fully green or, like the sectored parent, they had green and chlorotic sectors consistent with a heterozygous *sulf*/+ epigenotype with paramutation. The chlorotic sectors would have had the homozygous *sulf/sulf* epigenotype (referred to as *sulf*) and the non-paramutated green sectors would be *sulf* /+ (Fig. 1A).

**Figure 1:**
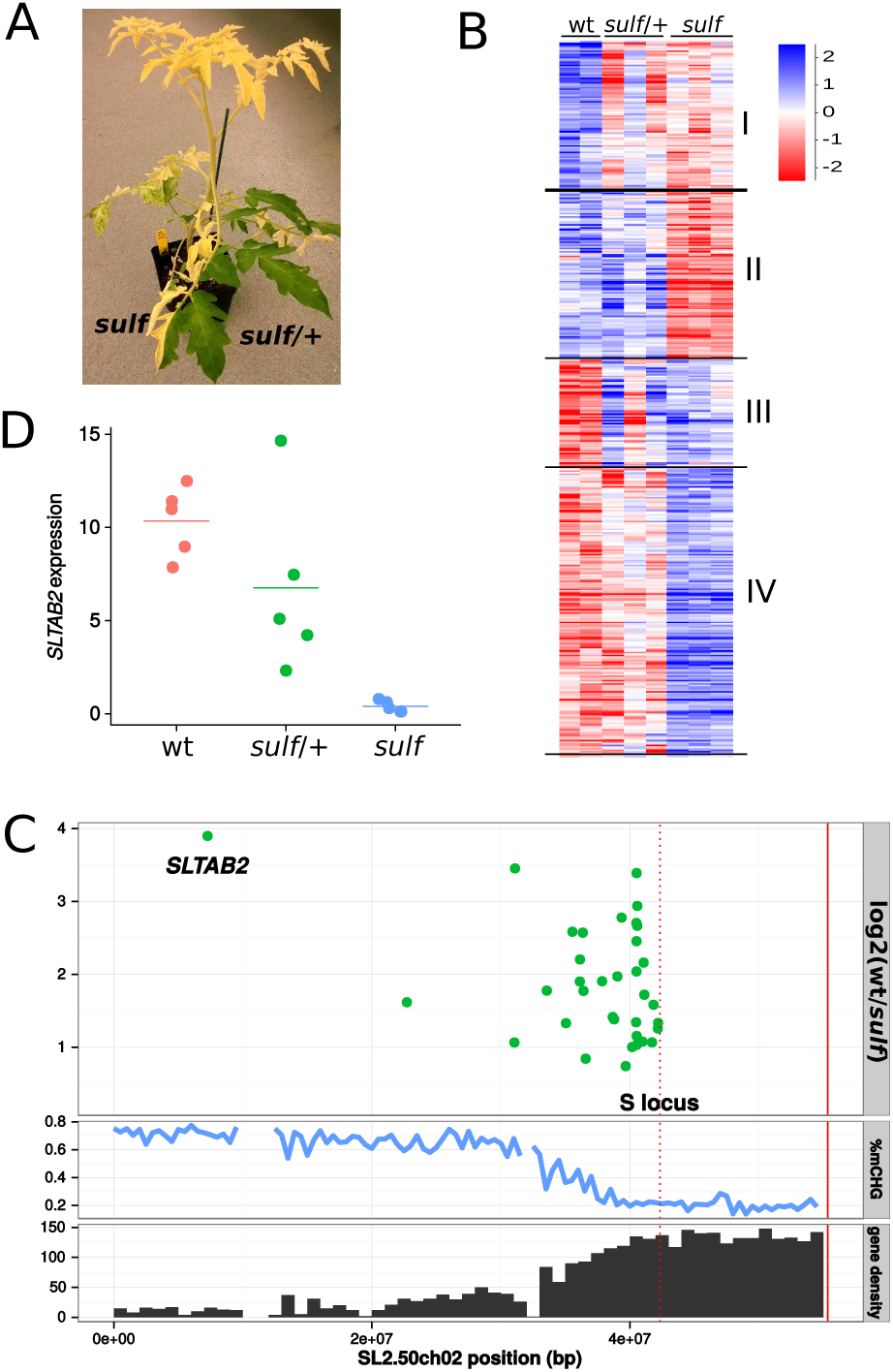
A. Fully paramutated yellow *sulf* tissue and non-paramutated green *sulf*/+ tissue on a plant grown from a heterozygous seed. B. Hierarchical clustering of 2237 differentially expressed genes between wt and *sulf*. With a correlation coefficient of 0.7, 2223 genes fall into 4 broad categories: down-regulated in *sulf* and sulf/+ (I), down-regulated in *sulf* only (II), up-regulated in *sulf* and *sulf*/+ (III), and up-regulated in *sulf* only (IV). Values are log2-transformed library-normalised/median-normalised counts. C. Down-regulated genes in *sulf* on chromosome 2. Log2 fold-change wt/sulf of significantly down-regulated genes (adjusted *p* < 0.05). Percentage CHG methylation and gene density (genes/Mb) are plotted along chromosome 2 to show the distribution of heterochromatin. The *S* locus (red dotted line) marks the right-most border for the possible location of *SULFUREA* (Hagemann and Snoad 1971). The end of the chromosome is marked by a solid red line. D. *SLTAB2* expression in wt, sulf/+ (green) and sulf (yellow) leaves. Horizontal bar shows the mean of 5 biological replicates. 26-fold reduction in sulf compared to wt (*pvalue* = 0.0002324, two-tailed t-test), while the difference between wt and sulf/+ is not significant (*pvalue* = 0.1771).

Based on the understanding of paramutation in maize, the *SULF* locus would be suppressed in *sulf* (paramutated yellow sectors), partially silent in *sulf* /+ (non-paramutated green sectors) and fully expressed in wild-type (wt). It would also be located in the pericentromeric heterochromatin of chromosome 2 upstream of the *S* locus (Hagemann and Snoad 1971). To find loci with these characteristics we analysed transcripts of wild-type, non paramutated *sulf*/+ and paramutated *sulf* leaves using mRNA-seq. We identified 2237 differentially expressed genes between *sulf* and wild-type (*p* < 0.05) that clustered into four main categories with distinct Gene Ontology enrichments (Fig. 1B and Table 1). Consistent with a decrease in photosystem I and the quantity of pigment in sulf leaves (Ehlert, Schottler, Tischendorf, Ludwig-Muller, and Bock 2008), many photosynthesis-related genes were down-regulated (Table 1, class II). The down-regulation of photosystem I was also detectable in the non-paramutated heterozygous *sulf*/+ leaves (Table 1, class I). Genes associated with various stress responses were up-regulated in *sulf* (Table 1, class IV) and were likely a secondary consequence of the *sulf* phenotype.

**Table 1:**
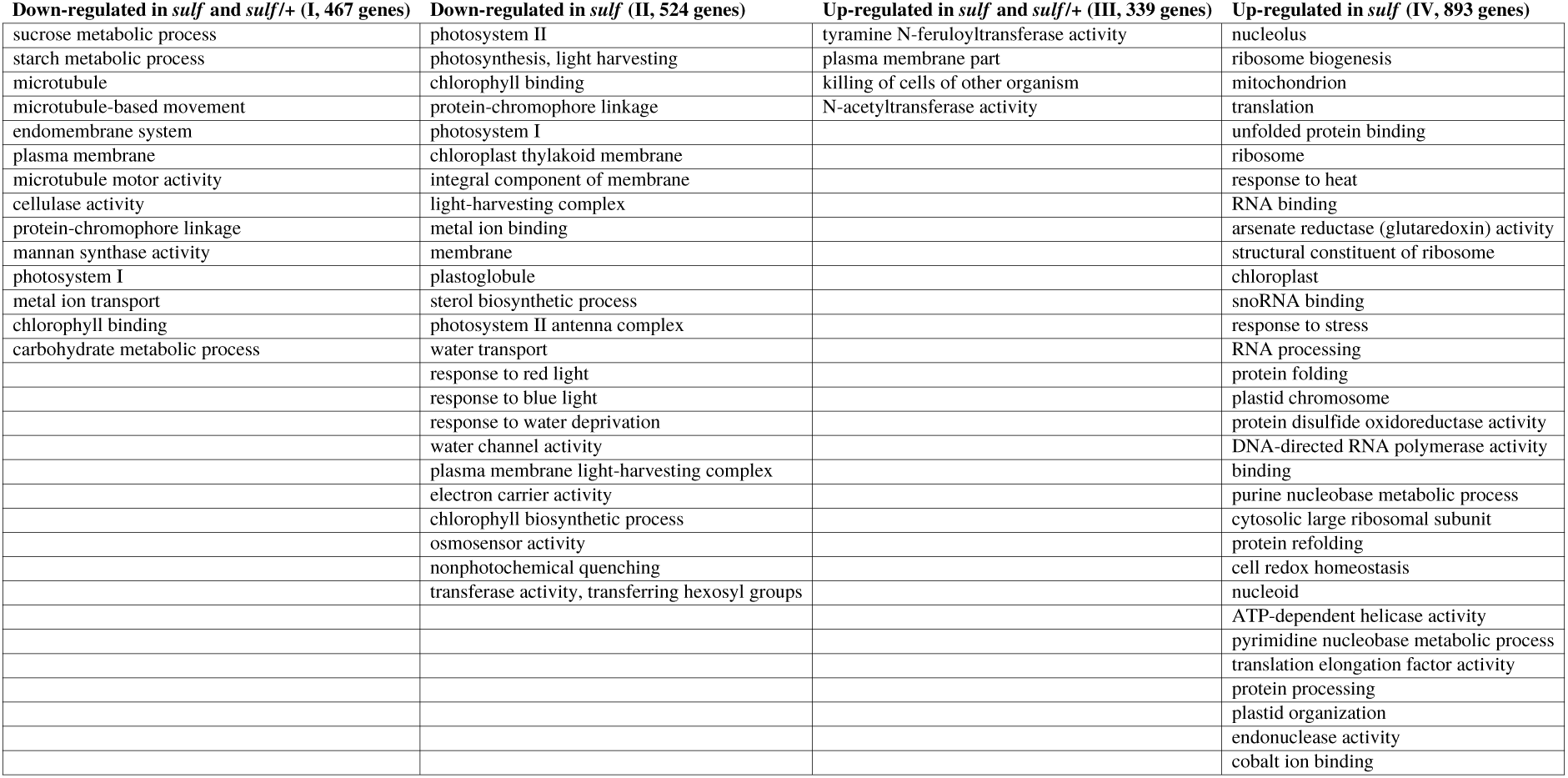
Enriched Gene ontology terms in the 4 hierarchical clusters

| Down-regulated in *sulf* and *sulf*/*+* (I, 467 genes) | Down-regulated in *sulf* (II, 524 genes) | Up-regulated in *sulf* and *sulf*/*+* (III, 339 genes) | Up-regulated in *sulf* (IV, 893 genes) |
| --- | --- | --- | --- |
| sucrose metabolic process | photosystem II | tyramine N-feruloyltransferase activity | nucleolus |
| starch metabolic process | photosynthesis, light harvesting | plasma membrane part | ribosome biogenesis |
| microtubule | chlorophyll binding | killing of cells of other organism | mitochondrion |
| microtubule-based movement | protein-chromophore linkage | N-acetyltransferase activity | translation |
| endomembrane system | photosystem I |  | unfolded protein binding |
| plasma membrane | chloroplast thylakoid membrane |  | ribosome |
| microtubule motor activity | integral component of membrane |  | response to heat |
| cellulase activity | light-harvesting complex |  | RNA binding |
| protein-chromophore linkage | metal ion binding |  | arsenate reductase (glutaredoxin) activity |
| mannan synthase activity | membrane |  | structural constituent of ribosome |
| photosystem I | plastoglobule |  | chloroplast |
| metal ion transport | sterol biosynthetic process |  | snoRNA binding |
| chlorophyll binding | photosystem II antenna complex |  | response to stress |
| carbohydrate metabolic process | water transport |  | RNA processing |
|  | response to red light |  | protein folding |
|  | response to blue light |  | plastid chromosome |
|  | response to water deprivation |  | protein disulfide oxidoreductase activity |
|  | water channel activity |  | DNA-directed RNA polymerase activity |
|  | plasma membrane light-harvesting complex |  | binding |
|  | electron carrier activity |  | purine nucleobase metabolic process |
|  | chlorophyll biosynthetic process |  | cytosolic large ribosomal subunit |
|  | osmosensor activity |  | protein refolding |
|  | nonphotochemical quenching |  | cell redox homeostasis |
|  | transferase activity, transferring hexosyl groups |  | nucleoid |
|  |  |  | ATP-dependent helicase activity |
|  |  |  | pyrimidine nucleobase metabolic process |
|  |  |  | translation elongation factor activity |
|  |  |  | protein processing |
|  |  |  | plastid organization |
|  |  |  | endonuclease activity |
|  |  |  | cobalt ion binding |

Of these differentially expressed genes, 36 were both down-regulated in *sulf* and located upstream of the *S* locus on chromosome 2. Among these candidates for *SULF*, Solyc02g005200 particularly stood out as being the most repressed in *sulf* (15-fold reduction) and at the predicted map position of *SULF* (29 cM from *S* locus, when the centromere-S distance is 30 cM). The other candidates mapped to the euchromatin or the transition zone between heterochromatin and euchromatin (Fig. 1C). Further qPCR analysis confirmed the strong down-regulation of Solyc02g005200 in *sulf* (26-fold) and revealed variable levels in *sulf*/+, compatible with mono-allelic expression (Fig. 1D). Solyc02g005200 is the ortholog of *Arabidopsis thaliana ATAB2* that is likely involved in the translation of mRNAs for both photosystems (Barneche, Winter, Crèvecœur, and Rochaix 2006) and we refer to it as *SLTAB2*.

To confirm the heritability of the *SLTAB2* silent epiallele we analysed *SLTAB2* in the F1 progeny of chlorotic *sulf* /+ (S. *lycopersicum* cv Lukullus) x *S. pimpinellifollium*. Out of 22 F1 plants, 8 displayed a paramutated phenotype with yellowing of parts of the leaves (Fig. 2A). These chlorotic plants expressed *SLTAB2* at half the level of wild-type plants (Fig. 2B) and, in a PCR test that differentiated the polymorphic alleles from the two parents, we only detected expression from *S. pimpinellifollium* (Fig. 2C). These data are consistent with *SLTAB2* being *SULFUREA:* in the green tissue the *S. pimpinellifollium* allele would have been expressed (but not the silent allele from the *sulf*/+ *S. lycopersicum* parent), and in the chlorotic tissue, it would have been paramutated. In addition, by confirming the heritable silencing of the *S. lycopersicum* allele in the chlorotic plants, these data confirm that *SLTAB2* silencing is not merely a consequence of the *sulf* phenotype.

**Figure 2:**
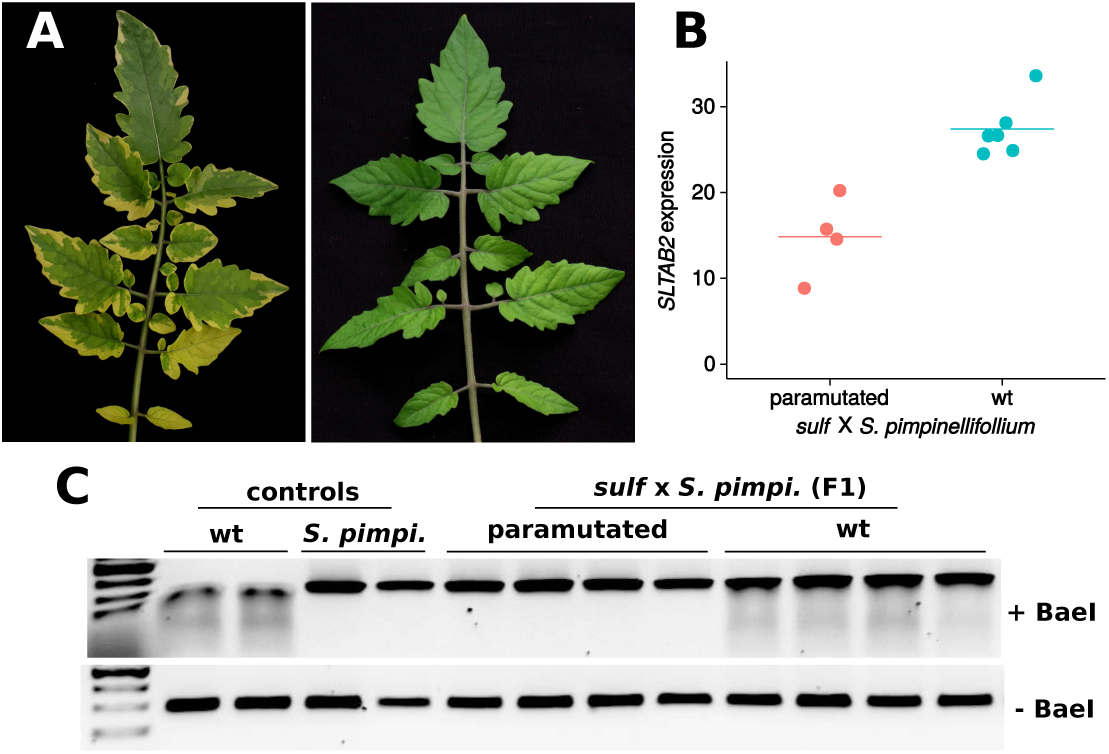
*SLTAB2* paramutation in the cross between a heterozygous *sulf* /+ and *S. pimpinellifollium*. A. Phenotypes of a paramutated (left) and wild-type leaf (right) in the F1. B. *SLTAB2* expression in the F1. Paramutated plants showed reduced expression (two-fold, *p* = 0.0057, two–tailed t-test). Mean represented by horizontal bar. C. Only the *S. pimpinellifollium SLTAB2* allele is expressed in paramutated F1 plants. The *S. lycopersicum* allele is sensitive to digestion by the BaeI restriction enzyme, resulting in a smear. Two SNPs in the *S. pimpinellifollium SLTAB2* allele make it resistant to BaeI treatment. F1s expressing the *S. lycopersicum* allele show a smear, whereas F1s in which the *S. lycopersicum* allele is silent do not.

Furthermore, consistent with equivalence of *SULF* and *SLTAB2*, the *Arabidopsis* T-DNA knockout of *ATAB2* is seedling lethal in heterotrophic conditions and deficient in green pigment ((Barneche, Winter, Crèvecœur, and Rochaix 2006) and Fig. 3A). This mutant also has less auxin than wild-type (Fig. 3B). These three phenotypes all resemble *sulf*.

**Figure 3:**
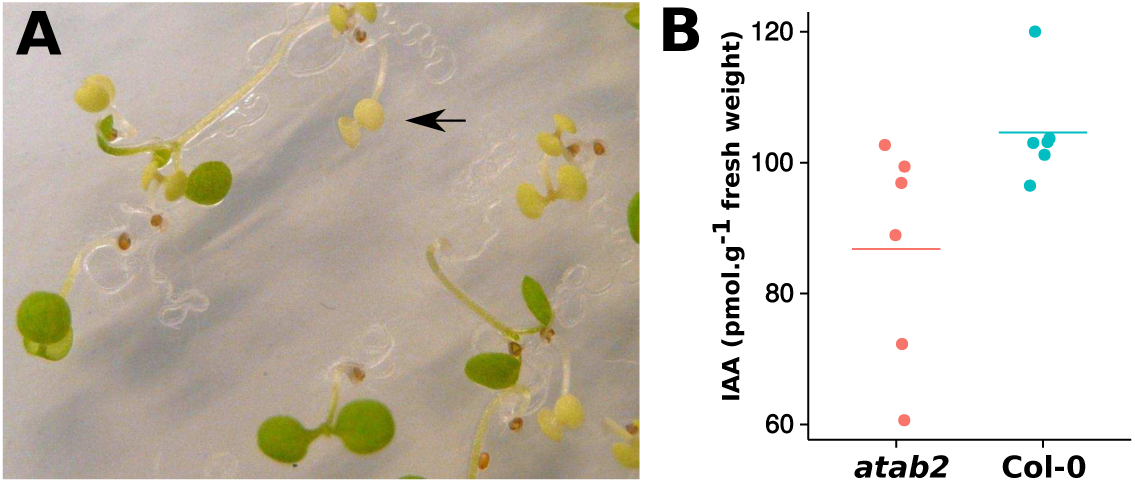
*Arabidopsis atab2* mutants resemble tomato *sulfurea*. A. Segregating *atab2* mutation in seedlings. Homozygous *atab2* mutants (arrow) are chlorotic and seedling lethal on heterotrophic medium. B. Decreased auxin in *atab2* (*p* − *value* = 0.02497, Kruskal-Wallis rank sum test).

### Paramutation is associated with changes in *SLTAB2* promoter DNA methylation

We predicted, based on the analysis of the maize *b1* gene (Stam, Belele, Ramakrishna, Dorweiler, Bennetzen, and Chandler 2002), that the DNA of the paramutagenic *sulf* would be hypermethylated. To find candidate loci based on this property we searched for differentially methylated regions (DMRs) based on the comparison of *sulf* and wild-type DNA in the appropriate region of chromosome 2 and adjacent to genes that were differentially expressed. Genome-wide there were thousands of such DMRs in the CHH context and hundreds in CG and CHG contexts with the CHH DMRs being predominantly hypermethylated in *sulf* whereas CG and CHG DMRs were evenly split between hyper- and hypo-DMRs (Fig. 4A). On chromosome 2 there were several differentially expressed genes with adjacent CHH DMRs but only the *SLTAB2* locus had strong DMRs in all cytosine contexts (Fig. 4B).

**Figure 4:**
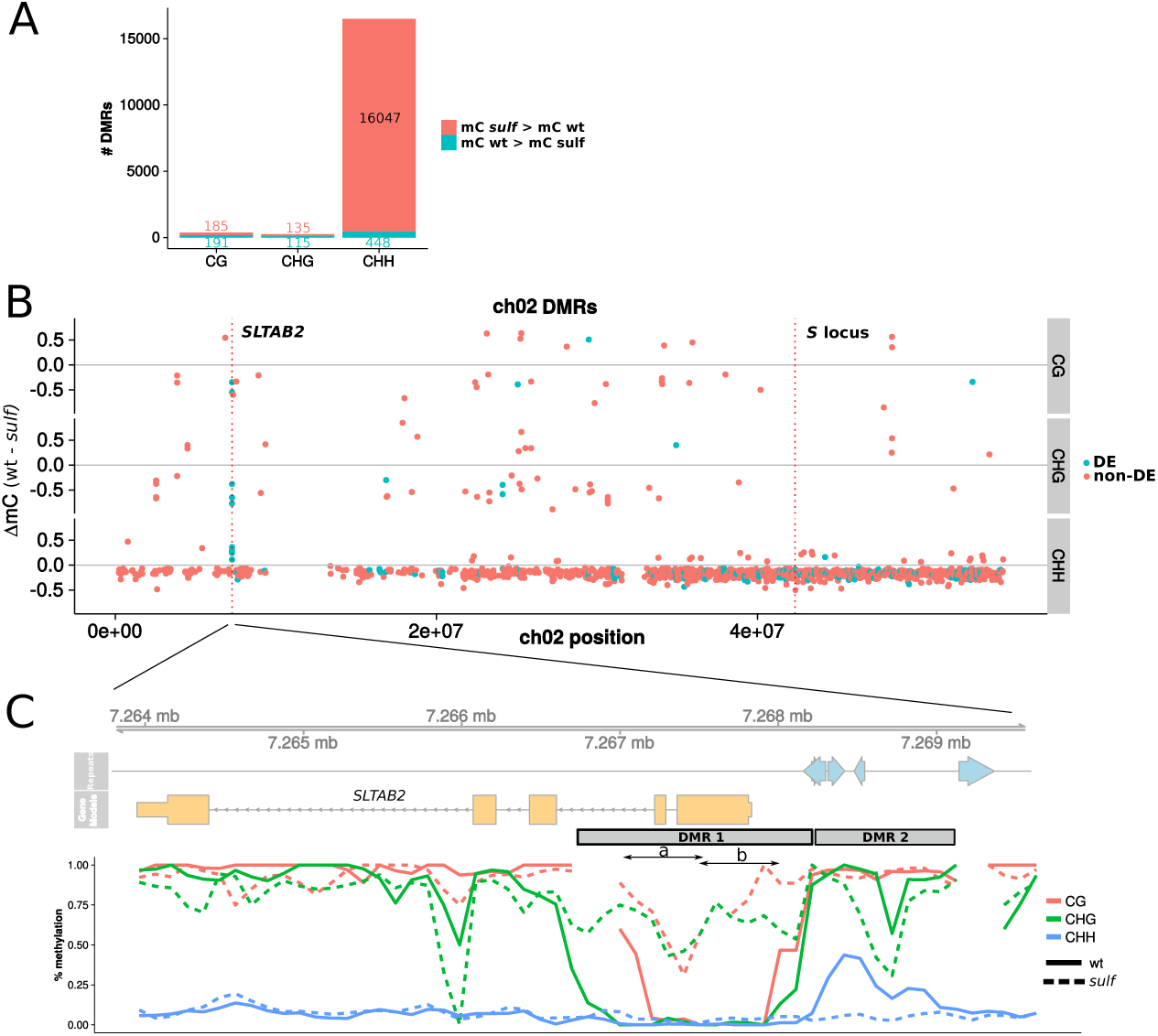
DMRs between wild-type and *sulfurea*. A. Hypo- and hyper-DMRs between wild-type and *sulfurea*. DMRs in the CHH context are the most abundant (1% of tested bins), and heavily biased for hypermethylation in *sulf*. B. Methylation difference (wt – *sulf*) of chromosome 2 DMRs. Negative values indicate hypermethylation in *sulfurea*, positive values hypomethylation. DMRs whose downstream gene is differentially expressed (DE) between wild-type and sulfurea are coloured in blue, while DMRs whose downstream gene is not differentially expressed (non-DE) are coloured red. C. Methylation over *SLTAB2* promoter, plotted in 200 bp sliding windows (step of 100 bp).

Closer inspection revealed that there are two adjacent DMRs in the immediate promoter of *SLTAB2*, DMR1 and DMR2 (Fig. 4C and Table 2). DMR2 overlaps annotated repeats directly upstream of *SLTAB2* and was hypomethylated in *sulf* in CHG and CHH contexts (78% and 4% methylation respectively, compared to 93% and 27% in wild-type). DMR1, in contrast, overlaps the transcriptional start site of *SLTAB2* from 300 bp upstream and encompassing the first two exons and introns, and it was hypermethylated in *sulf* in all contexts (73% mCG, 62% mCHG and 4% mCHH compared to 24%, 4% and 0.7% respectively in wild-type). This hypermethylation of the transcriptional start site of *SLTAB2* is consistent with a decrease in transcription in *sulf* and it further strengthens the case that *SLTAB2* is *SULF*.

**Table 2:** *SLTAB2* DMR methylation. Percentage methylation from two replicates. P-values from a logistic regression analysis on raw methylated and unmethylated cytosine counts.

| context | wt | *sulf* | p-value |
| --- | --- | --- | --- |
| DMR1 |  |  |  |
| CG | 21.4 ; 27.3 | 71.4 ; 76.8 | $< 2 * 10^{-16}$ |
| CHG | 2.3 ; 5.8 | 61.9 ; 62.4 | $< 2 * 10^{-16}$ |
| CHH | 0.5 ; 0.8 | 3.8 ; 4.8 | $1.12 * 10^{-7}$ |
| DMR2 |  |  |  |
| CG | 95.6 ; 96.4 | 96.6 ; 93.7 | 0.557 |
| CHG | 94.4 ; 90.9 | 78.0 ; 78.4 | $7.34 * 10^{-4}$ |
| CHH | 25.6 ; 29.3 | 5.5 ; 3.6 | $< 2 * 10^{-16}$ |

We also predicted based on the maize paramutation examples that *sulf* would correlate with 24-nt siRNAs. At a genome-wide scale, there was a distinct increase in 23–24-nt siRNAs in *sulf* compared to wild-type (Fig. 5A), in line with the pattern of CHH hypermethylation. At DMR1 of *SLTAB2*, the 23–24-nt siRNAs were more abundant in paramutated *sulf* (Fig. 5B) than in wild-type whereas at DMR2 the 23–24-nt siRNAs were less abundant than in wild-type (Fig. 5C).

**Figure 5:**
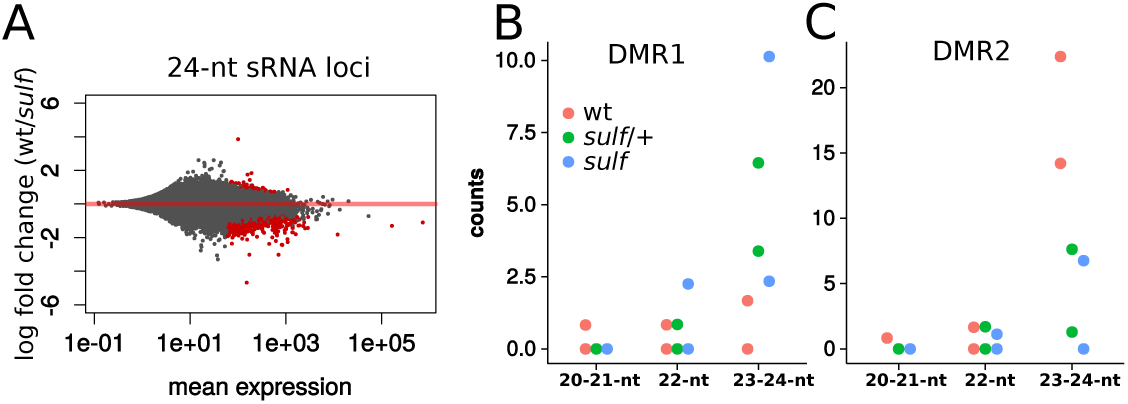
sRNAs in *sulfurea*. A. MA plot of 24-nt sRNA loci in wt and *sulf*. Of the differential sRNA loci between wt and *sulf* (in red, adjusted *p* < 0.05), 498 had more abundant sRNAs in *sulf*, while only 48 had more abundant sRNAs in wild-type. B. sRNA counts on DMR1 in wild-type, heterozygous *sulf* /+ and homozygous paramutated *sulf* leaves. 23–24-nt sRNAs are rare but more abundant in *sulf* (*p* = 0.0146, Poisson regression). C. sRNA counts on DMR2. 23–24-nt siRNAs are reduced in *sulf* (*p* = 5.48*e* − 05, Poisson regression). Counts in B and C are for normalised, uniquely mapping reads.

### VIGS of DMR1 durably silences *SLTAB2*

To further test the involvement of *SLTAB2* in *sulf* paramutation we used virus-induced gene silencing (VIGS). VIGS, when targeted to transcribed sequences, leads to knock-down of mRNA levels by post-transcriptional gene silencing but, when targeted at DNA sequences (e.g. *FWA* promoter in *A. thaliana* (Bond and Baulcombe 2015)), it can initiate heritable DNA methylation and transcriptional gene silencing. We cloned two segments (a and b, Fig. 4B) of DMR1 into tobacco rattle virus (TRV) RNA2 and inoculated them together with TRV RNA1 to wild-type tomato. While infection with TRV-DMR1a caused only mild variegation of the leaves, infection with TRV-DMR1b caused almost all plants to develop large *sulf*-like chlorotic sectors (Fig. 6A). The similarity of this VIGS phenotype to leaves of *sulf* is further evidence that *SLTAB2* and *SULF* are equivalent.

**Figure 6:**
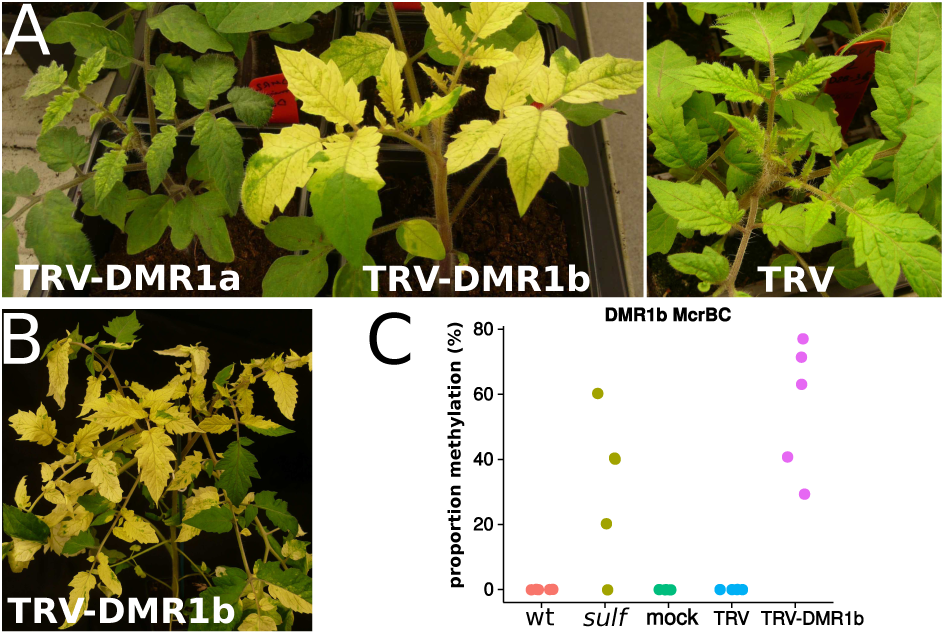
VIGS of *SLTAB2* DMR1. A. At 3 weeks post-infection, plants infected with TRV-DMR1b displayed large chlorotic sectors. B. These *sulfurea*-like sectors remained in later leaves at 2 months after infection. C. Methylation of DMR1 is detectable in chlorotic sectors.

DNA methylation analysis of chlorotic sectors by McrBC suggested that there is an epigenetic component to the silencing of *SLTAB2* by TRV-DMR1b : the targeted DNA was as strongly methylated as in *sulf* samples (Fig. 6C). The involvement of epigenetics is further supported by the lasting VIGS phenotype several months post-inoculation (Fig. 6B) during which time the level of the virus vector decreased. From these data we conclude that the DMR1b region of *SLTAB2* has the predicted characteristics of the paramutagenic component of *sulf* because it is susceptible to epigenetic modification.

## Discussion

A conclusive identification of *SLTAB2* as *SULF* would require the demonstration that an epigenetic modification is transferred from a silent allele in *sulf*/+ plants. The transfer would be to the previously active allele that, in turn, would transfer its modification in subsequent generations. It may be difficult to obtain such a demonstration because the *sulf/sulf* epigenotype is embryo lethal and it may not be possible to obtain seed in the plant sectors in which *sulf* /+ has converted to the *sulf/sulf* epigenotype.

The other evidence, however, is very strong that *SLTAB2* is *SULF:* it maps closer to *SULF* than any other genes with the predicted pattern of mRNA accumulation in *sulf* /+ and *sulf* tissue (Fig. 1B); it encodes a protein required for photosystem I production that explains the chlorotic phenotype; the orthologous *atab2* mutation has the same chlorosis and auxin deficient phenotype as *sulf* (Fig. 3); there is a definite epigenetic mark at the silent *SLTAB2* allele in *sulf* that is inherited from *sulf*/+ both in selfed and outcrossed progeny (Fig. 2); the epigenetic mark is associated with 24-nt siRNAs (Fig. 5) and, finally, VIGS targeted to the DMR can recapitulate both the physiological and epigenetic features of *sulf* (Fig. 6).

*SLTAB2* had been ruled out previously as *SULF* because it is expressed at detectable levels in *sulf* and it was thought that the mutation of the *Arabidopsis* orthologue could be rescued on sucrose (Ehlert, Schottler, Tischendorf, Ludwig-Muller, and Bock 2008). An alternative candidate for *SULF* was implicated in the tryptophan-independent pathway of auxin biosynthesis. With our new data, however, we show that the previous exclusion of *SLTAB2* was not valid because targeted suppression by VIGS or mutation of this gene produces an accurate phenocopy of the *sulf* phenotype including, in the *atab2* mutant, an auxin defect and seedling lethality.

A likely explanation for the *sulf* phenotype based on silencing of *SLTAB2* invokes the failure to translate the psaB mRNA as described for the orthologous mutations *ATAB2* in *Arabidopsis* and *TAB2* in *Chlamydomonas* (Barneche, Winter, Crèvecœur, and Rochaix 2006; Dauvillée, Stampacchia, Girard-Bascou, and Rochaix 2003). The PsaB protein is the reaction centre protein of photosystem I and, in its absence, the thylakoid membranes would fail to form, pigments would not accumulate at the normal levels and the leaves would be chlorotic. An auxin defect of *sulf* is a likely consequence of the PsaB defect, as observed in the *atab2* mutant (Fig. 3). In addition the genome-wide hypermethylation in the CHH context in *sulf* is reminiscent of transient hypermethylation in response to stress, already described in phosphate-starved rice (Secco, Wang, Shou, Schultz, Chiarenza, Ecker, Whelan, and Lister 2015) and virus-infected *Arabidopsis* (Bond and Baulcombe 2015).

The opportunity to study paramutation via *SLTAB/SULF* in tomato has several advantages over the various maize systems that have been most informative until now. First we have a VIGS system so that establishment of the epigenetic mark can be tracked directly in tomato mutants that are defective for components of the RNA silencing pathways. We will also be able to use VIGS on *sulf* /+ plants to test the role of various tomato genes in establishment and maintenance of paramutagenicity and paramutability.

A second benefit is the possibility to study the transfer of the epigenetic mark in vegetative tissue. With the well-studied maize paramutation systems this transfer is likely to occur early in embryo development and is not readily accessible to molecular analysis whereas, in tomato, it will be taking place in or close to vegetative meristems. It will still not be easy to access the cells in which the allelic transfer is taking place but we may be able to use the DMR1-specific siRNAs as markers of the transfer process. These RNAs are rare in total plant extracts (Fig. 5) but they may be more abundant at the primary sites of paramutation. Having identified the *SULF* gene we should also be able to complement the physiological consequences by providing a transgene without the target DNA of paramutation so that we can grow plants with a *sulf/sulf* epigenotype.

These various experimental tools will allow us to explore the differences of the tomato and maize paramutation systems. For example the apparent target of paramutation in *sulf* has no repeats and is close to the transcriptional start whereas in maize at the *B* locus they are essential and separated from the transcribed region by 100 kb. Answers to these and other questions will allow us to explore the frequency of paramutation-like events in plant breeding and evolution. Several recent findings indicate that such events are not restricted to the few well characterised examples of paramutation in maize and other species (Greaves, Groszmann, Wang, Peacock, and Dennis 2014; Regulski, Lu, Kendall, Donoghue, Reinders, Llaca, Deschamps, Smith, Levy, McCombie, Tingey, Rafalski, Hicks, Ware, and Martienssen 2013). They may be frequent and have an effect on transgressive and heterotic phenotypes.

## Acknowledgements

We thank Mel Steer and Jarmila Greplová for technical assistance, Donna Bond and Ottoline Leyser for advice and discussions, Fredy Barneche for sharing *atab2* seeds, and Adrian Valli, Jurek Paszkowski and Krys Kelly for critical reading of the manuscript. IPK Gatersleben kindly provided *sulfurea* and Lukullus seeds. This work was supported by the European Research Council Advanced Investigator Grant ERC-2013-AdG 340642, the Frank Smart Studentship and the Ministry of Education, Youth and Sports of the Czech Republic (the National Program for Sustainability I Nr. LO1204). D.C.B. is the Royal Society Edward Penley Abraham Research Professor.

## References

Axtell, M. J. 2013. “ShortStack: Comprehensive annotation and quantification of small RNA genes.” RNA 19: 740–751.

Barneche, Frédy, Veronika Winter, Michèle Crèvecœur, and Jean-David Rochaix. 2006. “ATAB2 is a novel factor in the signalling pathway of light-controlled synthesis of photosystem proteins.” The EMBO Journal 25: 5907–5918.

Bond, Donna M., and David C. Baulcombe. 2015. “Epigenetic transitions leading to heritable, RNA-mediated de novo silencing in *Arabidopsis thaliana*.” Proceedings of the National Academy of Sciences 112: 917–922.

Chandler, Vicki L., and Maike Stam. 2004. “Chromatin conversations: mechanisms and implications of paramutation.” Nature Reviews Genetics 5: 532–544.

Chitwood, D. H., J. N. Maloof, and N. R. Sinha. 2013. “Dynamic Transcriptomic Profiles between Tomato and a Wild Relative Reflect Distinct Developmental Architectures.” Plant Physiology 162: 537–552.

Coker, Jeffrey S., and Eric Davies. 2003. “Selection of candidate housekeeping controls in tomato plants using EST data.” BioTechniques 35: 740–748.

Dauvillée, David, Otello Stampacchia, Jacqueline Girard-Bascou, and Jean-David Rochaix. 2003. “Tab2 is a novel conserved RNA binding protein required for translation of the chloroplast psaB mRNA.” The EMBO journal 22: 6378–88.

Ehlert, B., M. A. Schottler, G. Tischendorf, J. Ludwig-Muller, and R. Bock. 2008. “The paramutated SULFUREA locus of tomato is involved in auxin biosynthesis.” Journal of Experimental Botany 59: 3635–3647.

Greaves, Ian K, Michael Groszmann, Aihua Wang, W James Peacock, and Elizabeth S Dennis. 2014. “Inheritance of Trans Chromosomal Methylation patterns from Arabidopsis F1 hybrids.” Proceedings of the National Academy of Science 111: 2017–22.

Hagemann, Rudolf. 1958. “Somatische Konversion bei Lycopersicon esculentum Mill.” Zeitschrift für Vererbungslehre 89: 587–613.

Hagemann, Rudolf. 1969. “Somatic Conversion (Paramutation) at the sulfurea Locus of Lycopersicon esculentum Mill. : IV. The Genotypic Determination of the Frequency of Conversion.” Theoretical and Applied Genetics 39: 295–305.

Hagemann, Rudolf and Brian Snoad. 1971. “Paramutation (somatic conversion) at the sulfurea locus of Lycopersicon esculentum. V. The localisation of sulf.” Heredity 27: 409–418.

Hollick, Jay B. 2012. “Paramutation: a trans-homolog interaction affecting heritable gene regulation.” Current Opinion in Plant Biology 15: 536–543.

Krueger, F. and S. R. Andrews. 2011. “Bismark: a flexible aligner and methylation caller for Bisulfite-Seq applications.” Bioinformatics 27: 1571–1572.

Liu, Y L, M Schiff, and S P Dinesh-Kumar. 2002. “Virus-induced gene silencing in tomato.” Plant Journal 31: 777–786.

Novák, Ondej, Eva Hényková, Ilkka Sairanen, Mariusz Kowalczyk, Tomáš Pospíšil, and Karin Ljung. 2012. “Tissue-specific profiling of the *Arabidopsis thaliana* auxin metabolome.” The Plant Journal 72: 523–536.

Regulski, Michael, Zhenyuan Lu, Jude Kendall, Mark T A Donoghue, Jon Reinders, Victor Llaca, Stephane Deschamps, Andrew Smith, Dan Levy, W Richard McCombie, Scott Tingey, Antoni Rafalski, James Hicks, Doreen Ware, and Robert A Martienssen. 2013. “The maize methylome influences mRNA splice sites and reveals widespread paramutation-like switches guided by small RNA.” Genome research 23: 1651–62.

Robinson, M. D., D. J. McCarthy, and G. K. Smyth. 2010. “edgeR: a Bioconductor package for differential expression analysis of digital gene expression data.” Bioinformatics 26: 139–140.

Secco, David, Chuang Wang, Huixia Shou, Matthew D Schultz, Serge Chiarenza, Joseph R Ecker, James Whelan, and Ryan Lister. 2015. “Stress induced gene expression drives transient DNA methylation changes at adjacent repetitive elements.” eLife p. 10.7554/eLife.09343.

Stam, Maike, Christiane Belele, Wusirika Ramakrishna, Jane E. Dorweiler, Jeffrey L. Bennetzen, and Vicki L. Chandler. 2002. “The regulatory regions required for B’ paramutation and expression are located far upstream of the maize b1 transcribed sequences.” Genetics 162: 917–930.

The Tomato Genome Consortium. 2012. “The tomato genome sequence provides insights into fleshy fruit evolution.” Nature 485: 635–641.

Urich, Mark A, Joseph R Nery, Ryan Lister, Robert J Schmitz, and Joseph R Ecker. 2015. “MethylC-seq library preparation for base-resolution whole-genome bisulfite sequencing.” Nature Protocols 10: 475–483.

